# The role of body temperature in regulating brain and body sizes in hominin evolution

**DOI:** 10.1101/2020.03.05.979351

**Authors:** Manasvi Lingam

## Abstract

A number of models have posited that the concomitant evolution of large brains and increased body sizes in hominins was constrained by metabolic costs. In such studies, the impact of body temperature has not been sufficiently addressed despite the well-established fact that the rates of most physiological processes are manifestly temperature-dependent. Hence, the role of body temperature in modulating the number of neurons and body size is investigated in this work by means of a simple quantitative model. It is determined that modest deviations in the body temperature (i.e., by a few degrees Celsius) might bring about substantive changes in brain and body parameters. In particular, a higher body temperature might prove amenable to an increase in the number of neurons, a higher brain-to-body mass ratio and fewer hours expended on feeding activities, while the converse applies when the temperature is lowered. It is therefore argued that future studies must endeavour to explore and incorporate the effects of body temperature in metabolic theories of hominin evolution, while also accounting for other factors such as foraging efficiency, diet and fire control in tandem.

## 1. Introduction

The human brain comprises the largest number of neurons among all primates and evinces a high degree of efficiency, economy and versatility [1]. The fact that humans have large brains, especially in relation to their body size, has been traditionally invoked to explain the emergence of their unique cognitive abilities [2, 3].^1^ In particular, the evolution of larger brains has been argued to confer a number of benefits ranging from the genesis, growth and navigation of complex cultural societies [7, 8, 9, 10, 11, 12, 13] to behavioural flexibility and general intelligence sensu lato [14, 15, 16, 17, 18, 19]. Although humans have the largest brains, they do not exhibit the largest body sizes of all primates. A number of explanations have been proposed for the relatively high brain-to-body mass ratio of Homo sapiens. One of the most well-known class of hypotheses posits that the concurrent evolution of large brains and bodies was inhibited due to the inherent metabolic costs [20, 21, 22]. In order to overcome the metabolic costs associated with large brains - which necessitate ∼ 20% of the total energy consumption in *H. sapiens* [23] - a number of avenues present themselves. Notable examples include: emergence of increasingly energy-efficient foraging strategies [24, 25, 26, 27, 28], shifts in the diet especially toward increased carnivory [29, 30, 31, 32, 33, 34], a reduction in the energy allocation to other organs (or functions) with inherently high energetic costs such as the gut [35, 36, 37] (see, however, Ref. [38]), and the adoption of cooking via fire control [39, 40, 41, 42, 43].

Thus, for a given energy intake by way of food intake, a tradeoff between the brain and body sizes is to be expected because this energy must be partitioned between the metabolic costs of sustaining the brain and the body. Interestingly enough, recent experiments have found support for an inverse correlation between brain development and the growth in body size from birth to adulthood in humans [44]. By drawing upon the notion of brain-body metabolic tradeoffs, Fonseca-Azevedo and Herculano-Houzel [45] developed a quantitative model to determine the constraints imposed on the sizes of hominin brains and bodies by metabolism. It was argued in Ref. [45] that the limited caloric yield of raw food substances was responsible for the relatively small brain sizes of great apes, and that a transition to cooked foods was potentially necessary to overcome this limitation in *Homo erectus*.

As noted earlier, other candidates for increasing the energy intake include the adoption of comparatively energy-rich diets, efficient foraging techniques, and cooking. Hence, it is conceivable that one (or more) of these methods played an essential role in relaxing the metabolic constraints on brain and body sizes [46]. There is, however, one crucial factor which has rarely been taken into account in current brain-body metabolic tradeoff models of hominin evolution, namely, the role of body temperature. This omission merits further scrutiny because there is ample empirical evidence for the sensitivity of numerous biological processes to the average body temperature, especially when it comes to the realm of metabolism [47, 48, 49].

As a consequence, the foremost aim of this work is to illustrate how small variations in the body temperature might translate to significant deviations in the brain and body sizes using a quantitative approach to draw general qualitative conclusions. To this effect, we will draw upon the methodology presented in Ref. [45] by constructing the simplest possible temperature-dependent model based on empirical and theoretical considerations. Along the way, we will also indicate what avenues must be pursued in the future to develop more sophisticated frameworks.

## 2 The mathematical framework

The details of the quantitative model are outlined, with a particular emphasis on the temperature-dependent aspects.

### 2.1 A primer on metabolic scaling

The centrality of metabolism in myriad biological processes is well-accepted [50, 48, 51], although other factors (e.g., hormones and environmental parameters) indubitably play a vital role [52]. A great deal of attention has been centered on uncovering the relationship between the basal metabolic rate (BMR) and the total mass (*M*) of the organism in question [53, 54]. Perhaps the best-known scaling law relating these two quantities is Kleiber’s Law [55, 56], which states that *R* ∝ *M* ^3/4^. The validity and universality of Kleiber’s Law remains the subject of extensive debate - although ongoing studies suggest that the 3/4 exponent is not universal across taxa [52, 57, 58, 59], it might nevertheless be reasonably accurate for larger mammals [60].

An oft-overlooked point, which forms the bedrock of our model, is that the BMR scales not only with the mass but also with the body temperature (*T*); this fact is along expected lines in view ofthe significance of temperature in biological functions [47, 49]. A widely used expression for the BMR (denoted by *R*) is

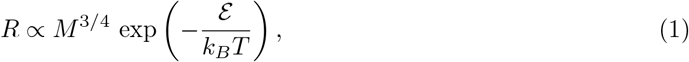

where *k_B_* is the Boltzmann constant and 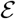 signifies the activation energy for the appropriate rate-limiting step in metabolism [61]. In other words, the above ansatz presumes that *R* is proportional to the Boltzmann-Arrhenius equation. As with Kleiber’s law, the effectiveness of this function across all taxa has generated debate [51, 60, 62], but we shall adopt this simple prescription because it constitutes a fairly robust leading-order approximation for unicellular organisms, plants and animals [58, 61, 63]. Across a wide spectrum of organisms, 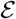 ranges between 0.6-0.7 eV in many (albeit not all) instances [48, 63, 64] and the mean value (0.65 eV) is nearly equal to the activation energy of 0.66 eV for ATP synthesis in mitochondria [65].

Let us adopt the temperature scaling in Eq. (1) and consider two organisms of the same mass, although at different body temperatures of *T*_0_ and *T*′ = *T*_0_ + Δ*T*. The ratio of the metabolic rates *R*(*T*′) and *R*(*T*_0_) (denoted by 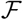) simplifies to

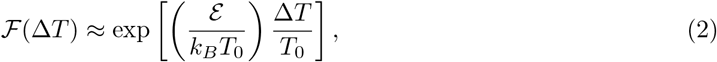

after invoking the ordering Δ*T*/*T*_0_ ≪ 1, which is valid for the range of Δ*T* considered in this work. To undertake our subsequent analysis, we will adopt 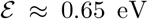 for reasons eludicated earlier. However, if we adopt the higher value 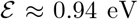 for mammals espoused in Ref. [66], our results remain qualitatively similar; in quantitative terms, Δ*T* must be replaced by 1.45Δ*T* instead. We shall also specify *T*_0_ ≈ 310 K (equivalent to *T*_0_ ≈ 37 °C) as it serves as the “canonical” temperature for present-day Homo sapiens. With these choices, Eq. (2) can be simplified further to yield

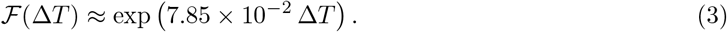

Note that Δ*T* can be either positive or negative in sign - owing to which we label it the residual temperature - implying that 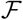 admits values both greater or smaller than unity.

### 2.2 The quantitative model

We are now in a position to ask the question. What would be the consequences for brain and body sizes if hominins such as *Homo erectus* possessed an ambient body temperature of *T*′ ≠ 37 °C? To answer this question, we will extend the model delineated in Ref. [45].

There are three major processes involved: (a) metabolic cost of body maintenance (*E*_BD_), (ii) metabolic cost of brain functioning (*E*_BR_), and (c) energetic intake via feeding (*E*_IN_). The expressions for each function are described below:

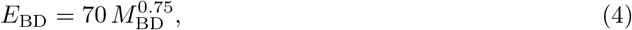

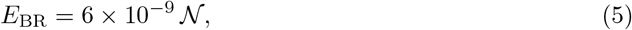

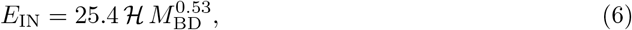

where the body mass MBD is measured in units of kg, 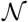 denotes the number of neurons, and 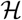 represents the number of hours (per day) spent on feeding activities. Note that all terms on the left-hand-side are measured in units of kCal per day.

It is apparent from inspecting Eq. (4) that this scaling is a consequence of Kleiber’s Law. However, as we have outlined earlier, the metabolic rate also exhibits a thermal dependence. Hence, we mustreplace *E*_BD_ with a modified metabolic cost that accounts for the body temperature. We implement this aspect by multiplying the right-hand-side of Eq. (4) with Eq. (3), as the latter embodies thermal deviations. Thus, the updated metabolic cost for body maintenance is expressible as

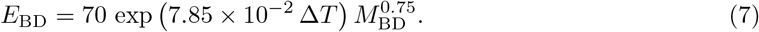

When it comes to energy expenditure by primates, the major component is basal metabolism [67]. We will therefore hold the prefactor of 70 in the above equation fixed, although variations of ≲ 50% are theoretically possible [46]. As the metabolic activity of the brain is intimately connected to the rest of the body, it is reasonable to assign a similar thermal scaling for *E*_BR_ [68], which leads us to

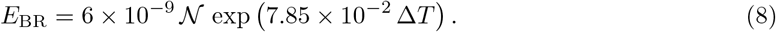

Next, let us turn our attention to Eq. (6). We can express the energy intake EIN as the product of the number of feeding hours per day 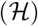 and the foraging efficiency 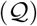; note that 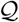 would have units of energy per unit time such as kCal/hour [46]. The allometric scaling exponent of 0.526 in Eq. (6) was derived empirically [45], but a theoretical explanation is feasible. We will assume that the foraging efficiency 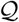 is proportional to the “search volume” (for resources) covered per unit time 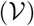. In turn, 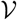 is modelled as being proportional to the rate of locomotion *υ* and the spatial reach *w*.

Thus, if we know the allometric scaling exponents *α* and *β* - where 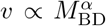 and 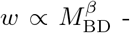 - we can determine 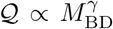 because our prior arguments indicate that *γ* ≈ *α* + *β*. As *w* should be governed by the length scale of the organism, we specify *β* = 1/3, while prior mathematical and empirical models for *υ* have yielded *α* ≈ 1/6-1/4 [69, 70]. Therefore, we obtain *γ* ≈ 0.5-0.583, which exhibits good agreement with the empirical scaling exponent of 0.526 in Eq. (6).

The last component we need to incorporate is the thermal dependence exhibited by *E*_IN_. For starters, we can assume that the energy input scales with the temperature as exp 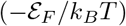, where 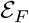 is the activation energy associated with feeding rates. By taking the ratio of the energy input rates at *T* and *T*_0_, we introduce the analog of (2), which is defined as

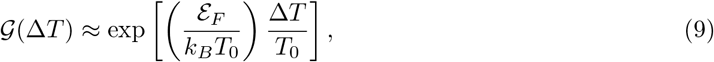

where 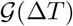 denotes the ratio. We will select 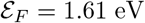, as this value was obtained for terrestrial endothermic vertebrates [71]; in the absence of primate-specific data, *faute de mieux*, this represents the best possible estimate. Upon substituting this value of 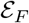 into the above expression and simplifying it, we end up with

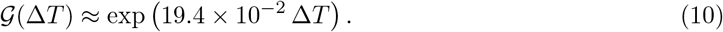

Therefore, the temperature-dependent version of *E*_IN_ is

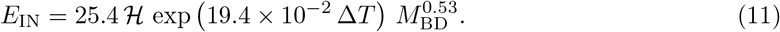

The appropriate criterion for facilitating the viability of a particular organism from a metabolic standpoint is *E*_IN_ > *E*_BD_ + *E*_BR_, where the functions are defined in Eq. (5), Eq. (6) and Eq. (7). The optimal limit is calculated by enforcing the following relation:

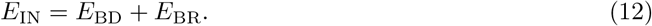

In turn, this imposes a constraint on MBD, 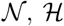 and Δ*T*. As the relationship between the former trio has already been investigated extensively in Ref. [45], we will focus primarily on Δ*T* herein.

## 3 Results

We commence our analysis by fixing 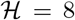 for the time being as it may constitute a sustainable upper bound on the number of hours devoted to feeding per day [45, 72]. For this choice of 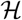, we can determine the compatible range of values admitted by *M*_BD_, 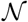 and Δ*T*. The results have been plotted in Fig. 1 for different choices of the residual temperature. We observe that there is a significant expansion in the number of permitted neurons (for a given body mass) if Δ*T* > 0 is adopted, and the converse trend is observed when Δ*T* < 0.

**Figure 1:**
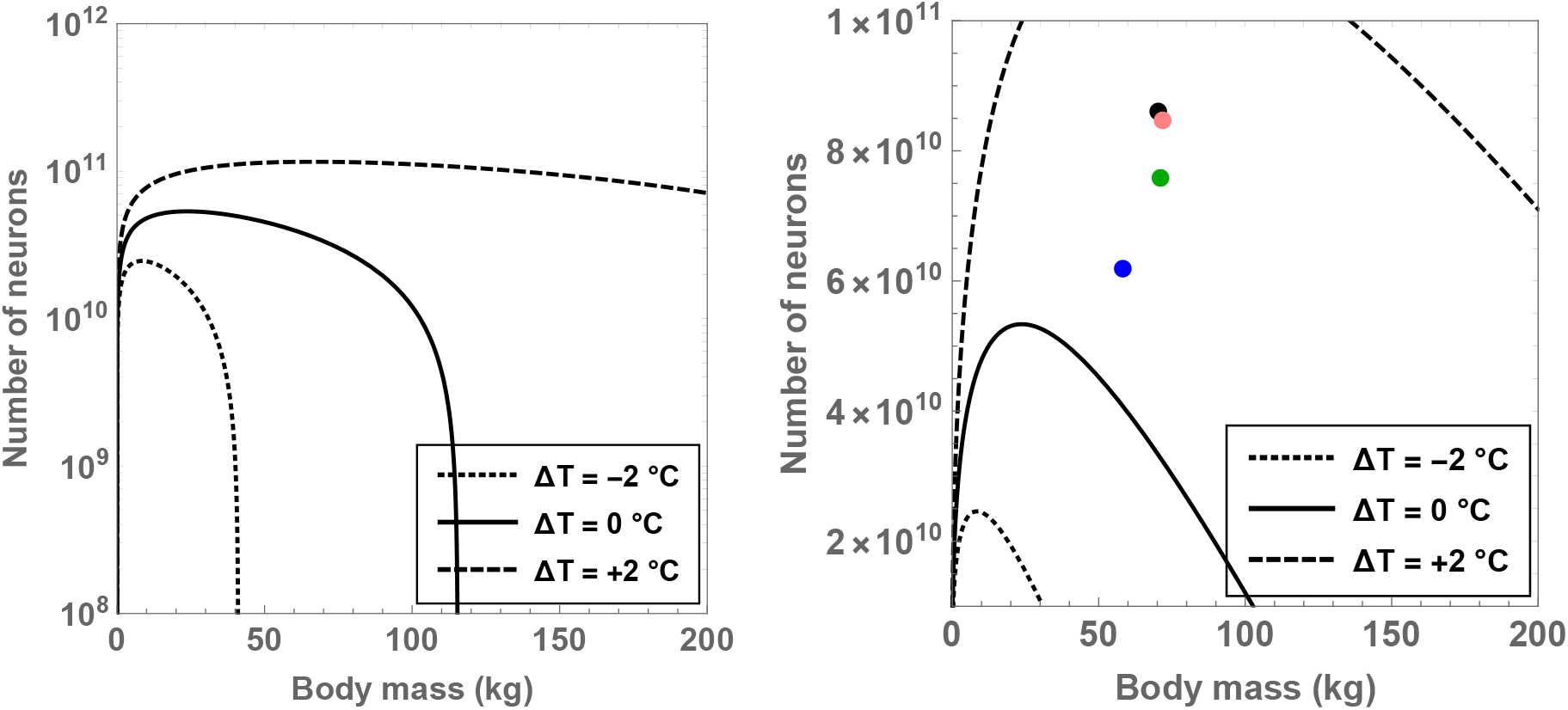
Compatible combinations of the number of neurons 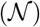 and body mass (*M*_BD_) are shown for putative primates that engage in 8 hours of feeding per day; the regions lying beneath the curves represent the permitted values. The dotted, unbroken and dashed curves correspond to body temperatures that differ from present-day humans by −2 ^°^C, 0 ^°^C and +2 ^°^C; the equivalent body temperatures are 35 ^°^C, 37 ^°^C and 39 ^°^C, respectively. In the right panel (which depicts a narrower range than the left panel), the black, orange, green and blue dots show the values of 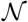 and *M*_BD_ for *H. sapiens, H. neanderthalensis, H. heidelbergensis* and *H. erectus*, respectively.

In particular, the right panel of Fig. 1 reveals that the existence of major hominins after *H. erectus* is not readily feasible if these species were characterized by a body temperature equal to (or lower than) present-day humans. On the other hand, in the scenario wherein these species possessed a body temperature higher than humans by 2 °C, their existence might be rendered mathematically viable. Although we have adopted the data for *M*_BD_ and 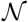 for various hominins from Table (S1) of Ref. [45], our results remain essentially unchanged even if we utilize more recent estimates for hominins [73].

In fact, we find that selecting 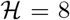 and Δ*T* = 1 °C in conjunction with the predicted mass of *H. erectus* yields 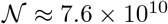, which is comfortably higher than the potential number of neurons characteristic of *H. erectus* 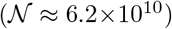 as per theoretical predictions [45]. Likewise, after choosing *M*_BD_ = 70 kg (motivated by *H. sapiens*), Δ*T* = 1.3 °C and 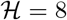, we arrive at 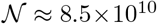 - a value that is close to the typical number of neurons in modern humans 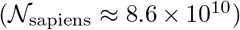. In contrast, upon adopting Δ*T* = 0 and holding all other parameters fixed, we end up with 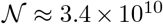, which is evidently much lower than the requisite 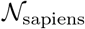.

Next, we turn our attention to the crucial brain-to-body mass ratio (denoted by *δ*_M_) and investigate the conditions necessary for fulfilling *δ*_M_ = 2%. As before, we shall restrict ourselves to 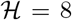 for reasons explained in the prior paragraph. To solve for *δ*_M_, we make use of the following relationship between *δ*_M_ and 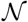 adopted from Ref. [74]:

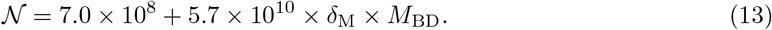

We can therefore obtain *δ*_M_ as a function of Δ*T* and *M*_BD_ for the given value of 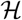. Upon inspecting Fig. 3 after doing so, the significance of the residual temperature in governing *δ*_M_ becomes manifest. For hominins with body temperatures equal to present-day humans, only modest masses of *M*_BD_ < 43 kg are permitted in order to ensure that *δ*_M_ > 2%. In contrast, if the body temperature is increased by 2 ^°^C, we find that the the maximal body mass that enables *δ*_M_ = 2% to hold true is *M*_BD_ ≈ 98 kg. As all documented hominins are less heavy than this critical value, it may have been feasible (from a mathematical standpoint) for these species to attain a high brain-to-body mass ratio provided that they had higher body temperatures. In fact, for 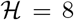 and Δ*T* = +1.5 ^°^C, we estimate that the maximum body mass compatible with *δ*_M_ = 2% is 80 kg, which is also larger than the typical masses of virtually all hominins.

Another result worth highlighting in this context concerns the maximum number of neurons for a given value of 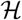 and Δ*T*. By solving for 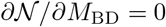, we end up with

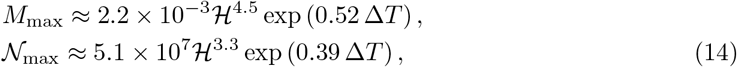

where 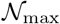 signifies the maximum number of neurons feasible and *M*_max_ represents the body mass of the hominin when 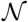 is attained. We have plotted *M*_max_ and 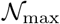 as a function of 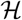 and Δ*T* in Fig. 2. We see that both of these functions are strongly dependent on the two variables. More specifically, we observe that *M*_max_ is close to the characteristic mass of *H*. sapiens when 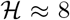 and Δ*T* = +2 °C; the corresponding value of the maximal neuron number is 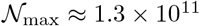, which is higher than 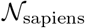. In contrast, if Δ*T* ≤ 0, we see that 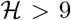 is imperative in order for 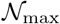 to overtake the characteristic number of neurons present in *H. sapiens*.

**Figure 2:**
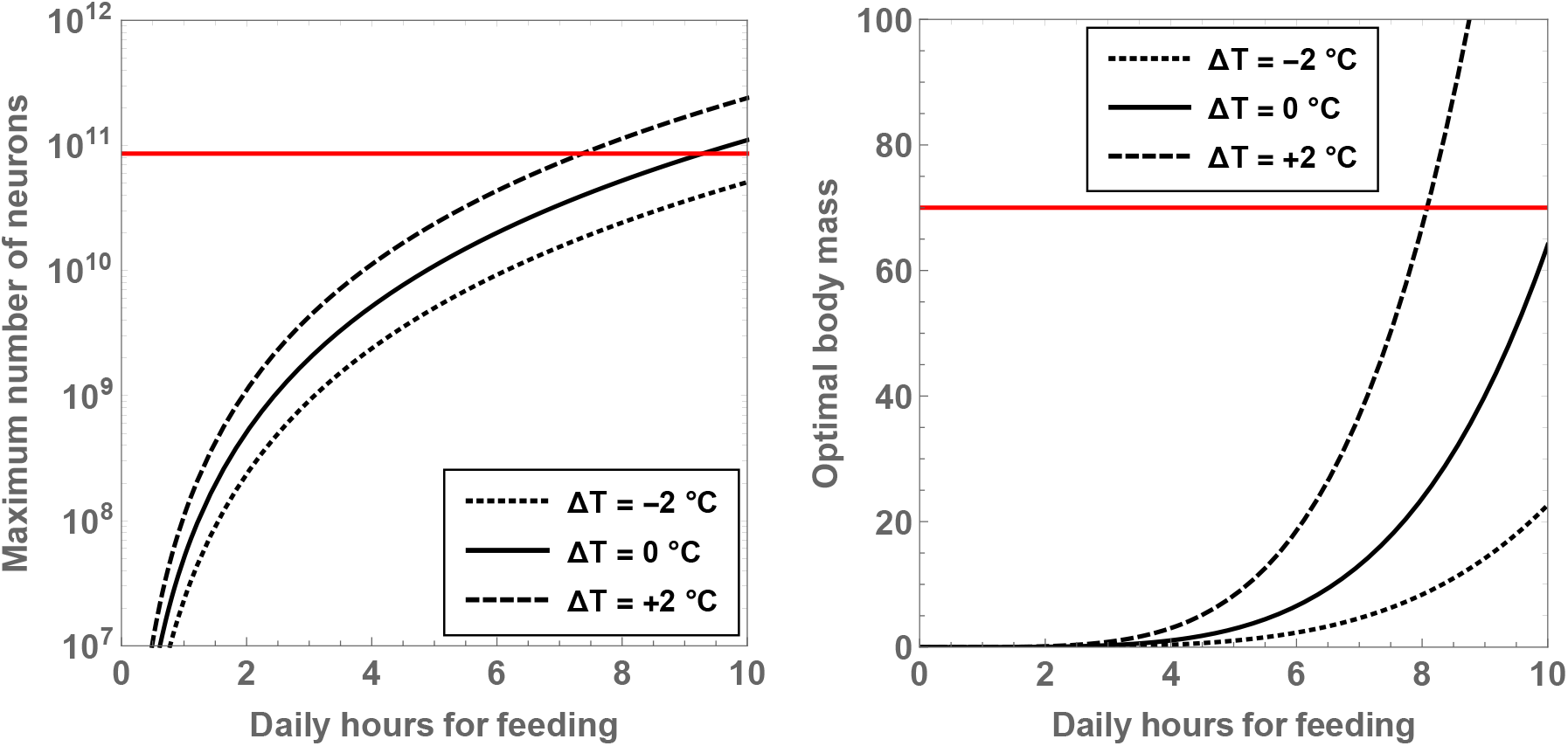
The maximal number of neurons (left panel) and the associated body mass (right panel) are shown as a function of the number of hours per day spent on feeding. The dotted, unbroken and dashed curves correspond to body temperatures that differ from present-day humans by −2 °C, 0 °C and +2 °C; the equivalent body temperatures are 35 °C, 37 °C and 39 °C, respectively. The horizontal red line depicts the neuron number (left panel) and mass (right panel) for *H. sapiens*.

**Figure 3:**
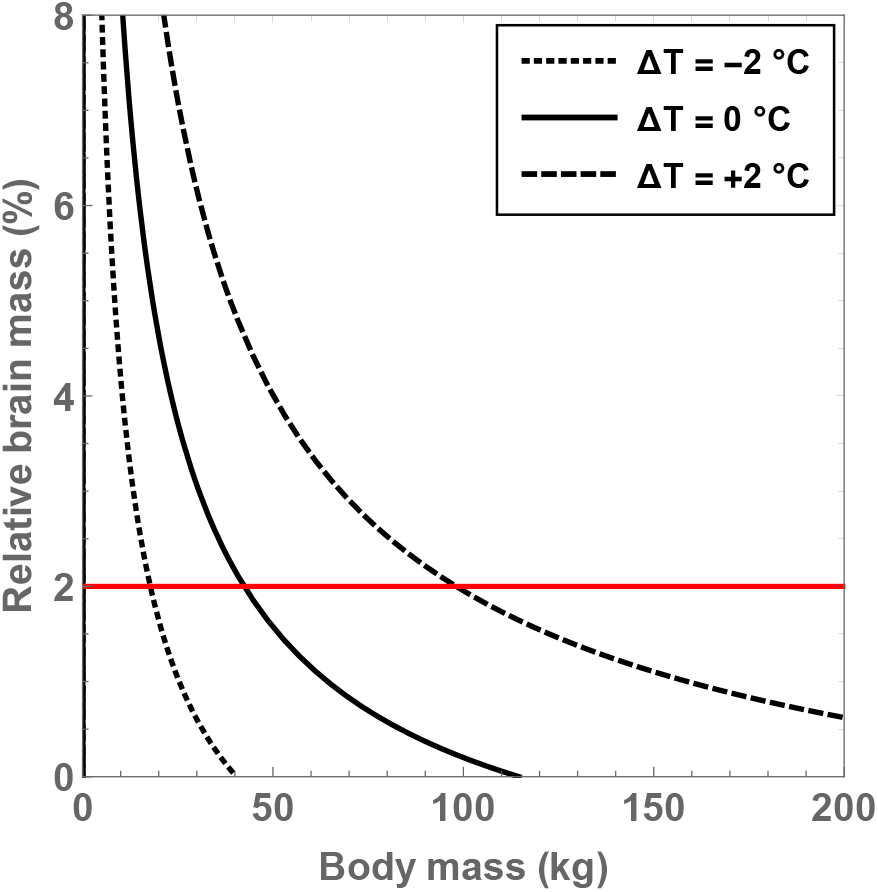
Compatible values of the brain-to-body mass ratio (*δ*_M_) and body mass (*M*_BD_) are depicted for putative hominins that take part in 8 hours of feeding per day; the regions within the curves are the permitted domains. The dotted, unbroken and dashed curves correspond to body temperatures that differ from present-day humans by −2 ^°^C, 0 ^°^C and +2 ^°^C; the equivalent body temperatures are 35 ^°^C, 37 ^°^C and 39 ^°^C, respectively. The horizontal red line is associated with *δ*_M_ = 2%.

Hitherto, we have focused on holding the number of feeding hours fixed and determining the constraints on the other variables. It is instructive to reverse the situation and hold 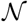 fixed, thereby allowing us to gauge the permitted range of 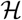, *M*_BD_ and Δ*T*. As the number of neurons presumably increased “only” by a factor of ~ 1.5 from *H. erectus* to *H. sapiens*, we can select a fiducial value of 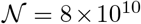 without much loss of generality; this number is close to the estimated number of neurons for *H. sapiens, H. neanderthalensis*, and *H. heidelbergensis* [1]. For this choice, the compatible values for the remaining variables have been plotted in Fig. 4. In common with the previous figures, we find that our results are sensitive to the temperature. In particular, it would seem impossible to achieve 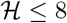 when the hominins in question have a temperature equal to (or lower than) present-day humans. In contrast, when we increase the body temperature by 2 °C, we find that 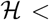 is achievable for 11 < *M*_BD_ < 182 kg.

**Figure 4:**
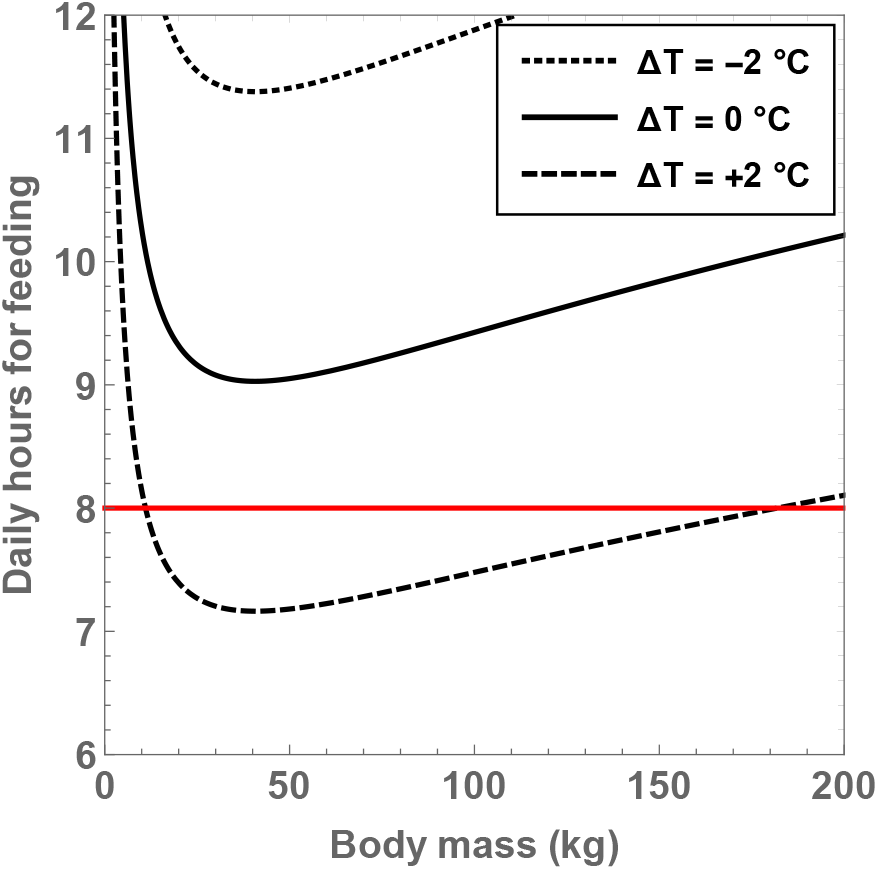
Compatible values of the number of hours per day expended on feeding 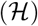 and body mass (*M*_BD_) are determined for putative hominins endowed with 80 billion neurons; the regions lying above the curves depict the allowed values. The dotted, unbroken and dashed curves correspond to body temperatures that differ from present-day humans by −2 °C, 0 °C and +2 °C; the equivalent body temperatures are 35 °C, 37 °C and 39 °C, respectively. The horizontal red line represents 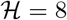 and can be regarded as a “sustainable” upper bound on 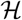.

By holding 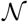 and Δ*T* fixed, we can determine the minimum number of hours that need to be expended on feeding as well as the associated body mass, which are denoted by 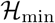 and *M*_min_, respectively. Thus, by solving for 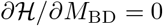, we arrive at

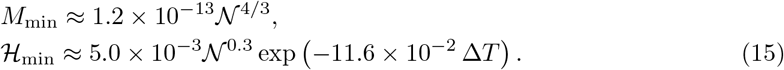

One of the striking aspects of this expression is that *M*_min_ does not exhibit any dependence on the temperature, although it does scale with the number of neurons. If we substitute the value of 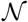 for *H. sapiens*, we find that the corresponding optimal body mass is 45 kg. Although this is approximately a factor of 1.5 removed from the typical mass of *H. sapiens*, it is important to appreciate that evolution is not necessarily guaranteed to converge toward strict optimality.

From the expression for 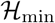, we see that the minimal number of feeding hours decreases with an increase of the residual temperature, which is consistent with our prior analysis. In Fig. 5, the behavior of 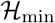 is shown as a function of 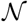 and Δ*T*. If the temperature of hominins is equal to (or lower than) the current body temperature of humans, we see that the minimal number of feeding hours becomes “large”, i.e., greater than 8 hours. For example, when we consider Δ*T* = 0 and 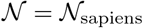, we arrive at the rather high value of 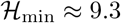. On the other hand, for Δ*T* = +2 °C and 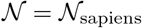, we obtain 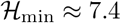, which is more reasonable.

**Figure 5:**
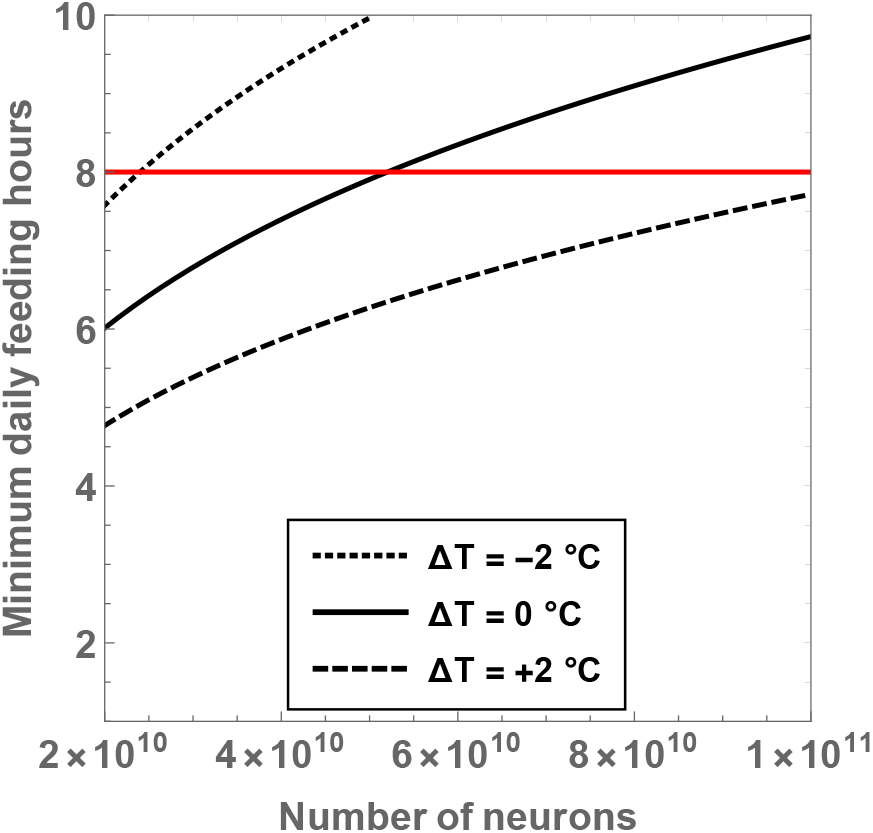
The minimum number of daily feeding hours is shown as a function of the total number of neurons. The dotted, unbroken and dashed curves correspond to body temperatures that differ from present-day humans by −2 ^°^C, 0 ^°^C and +2 ^°^C; the equivalent body temperatures are 35 ^°^C, 37 ^°^C and 39 ^°^C, respectively. The horizontal red line depicts the mass for *H. sapiens*.

Lastly, it is worth relaxing one of the key assumptions of Sec. 2 and exploring the ensuing consequences. Hitherto, we have implicitly utilized the “canonical” activation energy of 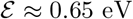 [48], but it has been suggested that adopting 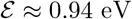 might be more accurate for mammals [66]. Upon recalculating 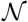 as a function of *M*_BD_ for this activation energy, we can evaluate the analogue of Fig. 1. The results are plotted in Fig. 6, from which it is apparent that a higher activation energy brings about smaller upward or downward shifts in 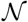 for a given value of Δ*T*.

**Figure 6:**
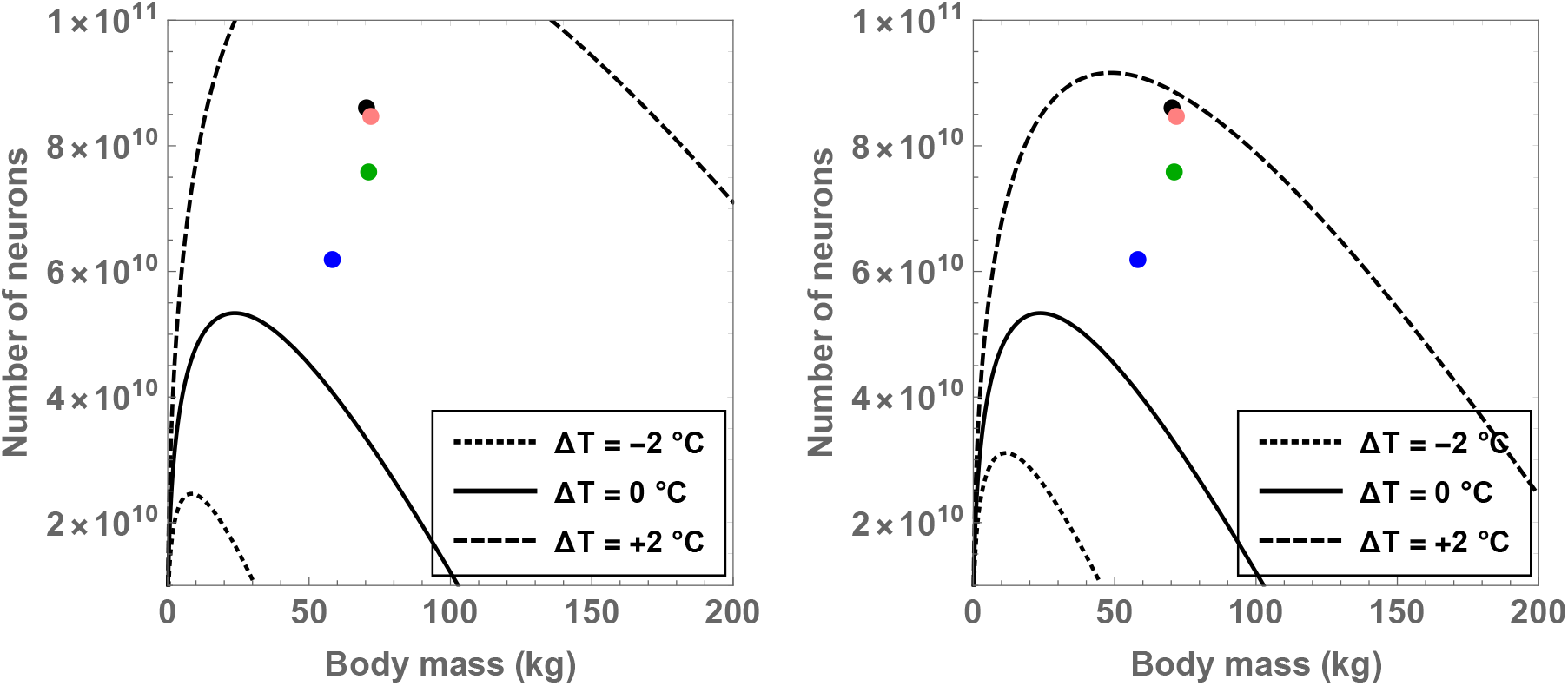
Compatible combinations of the number of neurons 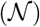 and body mass (*M*_BD_) are shown for putative primates that engage in 8 hours of feeding per day; regions below the curves represent the permitted values. The residual temperature (Δ*T*) quantifies the deviation of the body temperature from the conventional value of 37 °C. The left and right panels were derived using activation energies of 0.65 eV and 0.94 eV, respectively. The black, orange, green and blue dots show the values of 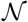 and *M*_BD_ for *H. sapiens, H. neanderthalensis, H. heidelbergensis* and *H. erectus*, respectively.

## 4 Conclusion

In this work, we explored the role of the body temperature in governing the tradeoff between body size and number of neurons (i.e., brain size) arising from metabolic constraints. There are two major qualitative conclusions that appear to be generally valid. First, even slight (i.e., a few °C) changes in the body temperature may engender substantial revisions of the permitted brain and body sizes. Second, we found that increasing the body temperature seem likely to drive putative hominins toward higher number of neurons, increased brain-to-body mass ratio and fewer hours spent on foraging to cover metabolic needs. The converse effects are ostensibly true when the body temperature is raised. From a quantitative standpoint, we estimated that a body temperature which is approximately 1-2 °C higher (at the minimum) than *T*_0_ ≈ 37 °C might suffice to offset metabolic costs and permit the evolution of hominins starting with *H. erectus*; on the other hand, ceteris paribus, the canonical human body temperature would pose difficulties for their emergence.

The next question that arises is whether a higher body temperature was feasible in the past. A very recent study concluded that the body temperature of humans has decreased (in a linear fashion) by 0.6 °C in the past ~ 150 years [75]. The fact that this marked change in the temperature purportedly occurred over a relatively short timescale suggests that hominins may have possessed higher body temperatures with respect to present-day humans. Note that the emergence of relatively large-brained hominins commencing with *H. erectus* took place during the interval of ~ 0.2-2 Mya [76, 77, 78]. Hence, the requisite average increase in the body temperature is ~ 10^−5^ to ~ 10^−6^ °C per year under the supposition of linear scaling to achieve a cumulative alteration of the body temperature by Δ*T* = 2 °C over the chosen timescale. In turn, the linear model translates to a fairly minimal increase of ~ 10^−3^ − 10^−4^ °C per century; whether this is sustainable over a long timescale is certainly debatable, but it cannot be dismissed *a priori*.

We can also ask ourselves the converse question of what would happen if lower body temperatures were prevalent. Before addressing this issue, it is worth examining the thermal data from primates. The body temperature of the western lowland gorilla (*G. g. gorilla*) is around 35.5 °C [79], which is approximately 1.5 °C lower than the fiducial value employed in our analysis. On the other hand, both chimpanzees (*P. troglodytes*) and Bornean orangutans (*P. pygmaeus*) have body temperatures nearly equal to that of humans [80]. Several species of *Euarchonta* (which encompasses primates) evince body temperatures < 35 °C [60]. Thus, we cannot rule out the possibility that hominins had lower body temperatures, especially given that dietary restrictions have been documented to play a vital role in regulating human body temperature [81].

In the event that Δ*T* < 0, i.e., hominins had lower body temperatures, the metabolic tradeoffs are rendered far more stringent. Broadly speaking, as per the results presented earlier, one can expect fewer number of neurons, decreased brain-to-body mass ratio and more hours spent on foraging to cover metabolic needs. In other words, the necessity for a counteracting mechanism also increases commensurately. An evident solution is to enhance the energetic intake per hour of feeding. This outcome can be achieved by changes in diet and foraging [29, 82, 32], or through the consumption of cooked foods [83, 42]. Thus, in case putative hominins were characterized by lower body temperatures, the “cooked food” hypothesis pioneered by Richard Wrangham and collaborators [39, 40, 43] acquires greater significance. While we do not have an unambiguous picture of whether genus Homo was capable of fire control in the early and mid Pleistocene, recent developments on this front - especially in connection with the site FxJj20 in Koobi Fora, Kenya [84, 85] - seem promising [86].

To sum up, if the body temperature of hominins was higher or lower than current humans by a few °C, our analysis indicates that this shift potentially engendered substantive changes in their body sizes and number of neurons insofar as metabolic constraints are concerned. Therefore, at the minimum, there are two interconnected areas that merit further investigation: (i) accurately estimating hominin body temperature by synthesizing empirical and theoretical studies in physiology, anthropology, palaeontology and genomics, and (ii) utilizing the inferred temperature in sophisticated thermal models (e.g., with quadratic corrections) to comprehend how metabolic (*E*_BD_ and *E*_BR_) and feeding (*E*_IN_) costs and the resultant brain and body sizes are modulated. Needless to say, aside from the role of temperature, a multitude of extrinsic and intrinsic factors such as the foraging efficiency, cumulative culture, diet, and climate must also be taken into account.

## Acknowledgments

It is a pleasure to thank Richard Wrangham for the encouraging and insightful feedback regarding the paper.

## Footnotes

1 It is, however, important to recognize that large brains are not necessarily better due to the fact that additional factors such as “modularity and interconnectivity” [4] also determine several aspects of cognition [5, 6].

